# Helping behavior is associated with increased affiliative behavior, activation of the prosocial brain network and elevated oxytocin receptor expression in the nucleus accumbens

**DOI:** 10.1101/2024.05.06.592793

**Authors:** R. Hazani, J.M. Breton, E. Trachtenberg, B. Kantor, A. Maman, E. Bigelman, S. Cole, A. Weller, I. Ben-Ami Bartal

**Affiliations:** Psychology Department, Bar-Ilan University, Ramat Gan, Israel, 5290002; Gonda Brain Research Center, Bar-Ilan University, Ramat Gan, Israel, 5290002; Geha Mental Health Center, Petah Tikva, Israel, 4917002; Department of Psychology, Northeastern University, Boston, MA, 02115, USA; School of Psychological Sciences, Tel-Aviv University, Tel Aviv, Israel, 6997801; Sagol School of Neuroscience, Tel-Aviv University, Tel Aviv, Israel, 6997801; UCLA School of Medicine, Los Angeles, CA, 90095, USA

**Author notes:** equal contribution (R. Hazani, J.M. Breton). Corresponding Authors and email addresses: Correspondence should be addressed to either Jocelyn Breton at or Inbal Ben-Ami Bartal at.

**Keywords:** prosocial, social affiliation, oxytocin receptor, nucleus accumbens, neural network

## Abstract

A prosocial response to others in distress is increasingly recognized as a natural behavior for many social species, from humans to rodents. While prosocial behavior is more frequently observed towards familiar conspecifics, even within the same social context some individuals are more prone to help than others. For instance, in a rat helping behavior test, rats can release a distressed conspecific trapped inside a restrainer by opening the restrainer door. Typically, rats are motivated to release a trapped cagemate, and consistently release the trapped rat (‘openers’), yet around 30% do not open the restrainer (‘non-openers’). To characterize the difference between these populations, behavioral and neural activity were compared between opener and non-opener rats tested with a trapped cagemate in the helping test. Behaviorally, openers showed significantly more social affiliative behavior both before and after door-opening compared to non-openers. Analysis of brain-wide neural activity based on the immediate early gene c-Fos revealed increased activity in openers in the previously identified prosocial neural network compared to non-openers. The network includes regions associated with empathy in humans (somatosensory cortex, insula, cingulate cortex and frontal cortex), and motivation and reward regions such as the nucleus accumbens. Oxytocin receptor mRNA expression levels were higher in the accumbens but not the anterior insula. Several transcription control pathways were also significantly upregulated in openers’ accumbens. These findings indicate that prosocial behavior may be predicted by affiliative behavior and activity in the prosocial neural network and provide targets for the investigation of causal mechanisms underlying prosocial behavior.

**Significance Statement:** Prosocial behavior is observed in many social species, including rodents, yet the determinants underlying why some animals help and others do not is poorly understood. Here, we show behavioral and neural differences between prosocial and non-prosocial pairs in a rat helping behavior test, with increased social interaction and nucleus accumbens oxytocin receptor gene expression in animals that helped.

## Introduction

The motivation to help distressed others has been increasingly demonstrated across social species (Rault, 2019; Wu and Hong, 2022), extending beyond parental care and bonded pairs to conspecifics of the same social group (Decety et al., 2016). The neural mechanisms underlying a prosocial response involve processing cues of distress in others, which may elicit empathic arousal in the observer and motivate acts meant to improve the others’ well-being, such as consolation or targeted helping. The response to a distressed conspecific is critically different than social interaction in neutral contexts, and in many species (rodents, primates, elephants, corvids) elicits approach and consolatory touch (de Waal and Preston, 2017), considered effective in reducing distress. In recent years, more sophisticated prosocial acts have been demonstrated, which require actions congruent with the conspecific’s goals such as food sharing (Hernandez-Lallement et al., 2014; Marquez et al., 2015; Brucks and von Bayern, 2020; Nafcha et al., 2023), and rescue behaviors (Moscovice et al., 2004; Ben-Ami Bartal et al., 2011; Sato et al., 2015; Zhang et al., 2024). These behaviors are preferentially demonstrated for affiliated others, either on the individual or group level. However, in all these experiments, some proportion of subjects fail to show prosocial behavior, with reports ranging between 30%-50%. These individual differences could be explained by reduced sociability or prosocial motivation in non-helpers, failure to learn the task, high levels of personal distress, low trait empathy, or specific social dynamics. In this study, we aimed to understand determinants of prosocial behavior by outlining the behavioral and neural differences between prosocial and non-prosocial pairs in a rat helping behavior test (HBT), whereby rats can help a trapped conspecific by opening a restrainer door, freeing the trapped rat. In the HBT, ∼70% of rats typically release a trapped cagemate, and once they learn to open the restrainer door, they tend to repeat the behavior quickly and consistently on following sessions (Ben-Ami Bartal et al., 2011). Helping depends on the transfer of distress between the free and trapped rat (Ben-Ami Bartal et al., 2016) and moreover, rats selectively help affiliated others, not releasing trapped strangers of an unfamiliar strain (Ben-Ami Bartal et al., 2014). The role of distress in motivating helping, combined with the social selectivity of the behavior, are suggestive of empathic processes, where the affective state of one individual induces a congruent state in the observer, coupled with prosocial motivation to act for their wellbeing.

The neural network recruited during the HBT involves central regions of the human empathy network, including the anterior cingulate cortex (ACC), anterior insula (AI), and medial prefrontal cortex (mPFC), among others. In addition, activity in the reward network was correlated with helping (Ben-Ami Bartal et al., 2021). The nucleus accumbens (NAc) is a main hub of this “prosocial brain network”, with increased NAc activity observed in the presence of trapped ingroup members (cagemates or strangers of the same strain) compared to outgroup members (strangers of an unfamiliar strain) who weren’t helped (Ben-Ami Bartal et al., 2021). Increased NAc activity was also found in adolescent rats who released outgroup members, compared to adult rats who did not (Breton et al., 2022). This pattern of neural activity supports the idea that rats process the distress of the trapped rat and that an interaction between affective arousal and social motivation is required for helping to occur.

Here, the brain-wide response during the HBT was compared between helper rats, who consistently released trapped cagemates (“openers”) and rats who did not release cagemates (“non-openers”). We hypothesized that if non-openers were characterized by reduced prosocial motivation, we would observe reduced response in the prosocial brain network described above. To investigate whether oxytocin (OXT) signaling could account for differences in prosociality, OXT receptor mRNA expression levels were measured in the NAc and AI via RNAseq and qPCR. Behavioral metrics of social affiliation were measured to assesses whether social relationship strength was predictive of future helping.

## Materials and Methods

### Experimental design

#### Animals

Experiment 1 included a total of 32 male and 32 female adult Wistar rats (Envigo RMS, Israel), and was performed at Bar Ilan University. The experiment was performed in three batches to ensure consistency with replication. All rats arrived at the animal facility on postnatal day (PND) 52 and acclimated to the facility and their cagemate for 2.5 weeks before behavioral testing began. They were housed in same sex pairs and received water and food ad libitum. The room had a controlled temperature of 22±2 °C and a 12-hour light/dark cycle (lights on at 07:00 AM). At the beginning of the test phase, the rats’ weights were 192-245 g for females and 310-423 for males, and weights were recorded once a week throughout the experiment. The study protocol conformed to Society for Neuroscience guidelines and was approved by the Institutional Animal Use and Care Committee at Bar Ilan University.

Experiment 2 included 13 adult male Sprague-Dawley (SD) rats, and was performed at the University of California, Berkeley. Data from these animals has been previously published using different analyses, and full details can be found in prior work (see Ben-Ami Bartal et al., 2021). This experiment was performed in accordance with protocols approved by the Institutional Animal Care and Use Committee at the University of California, Berkeley. In both experiments, every effort was made to minimize animal suffering and to reduce the number of animals used.

#### Behavioral Testing

All procedures were undertaken during daylight hours. Throughout testing days, the pairs were tested in a counterbalanced order to ensure that testing order did not bias their behavior. The testing arena was made of white Polygal (50 w X 50 d X 70 h cm). All behavioral apparatuses were sanitized with a 70% ethanol solution at the end of each test period to remove odor residue. Tests were filmed using a video camera connected to Ethovision XT 15 software (Noldus, Netherlands). Manual behavior analysis was coded using the Solomon Coder software (Péter, 2011).

##### Habituation

After acclimation, rats underwent five days of habituation, during which they were accustomed to the experimenters, the experimental room, and the testing arena (**Fig. 1A**). This procedure was carried out to ensure that the only novel and stressful element during the HBT was the presence of a rat trapped inside the restrainer. On the first day, the rats were transported with their home cages to the testing room and remained there for 15 minutes without interruption before undergoing the Boldness Test. On days 2-5, after the Boldness Test, the experimenters handled the rats for 5-minutes. Afterward, each pair was placed in the testing arena together (without the restrainer) for 30-minutes. Social Interaction Test (SIT) measures were obtained on day 2 of habituation. On the 5th day, after the handling, the Open Field Test (OFT) was conducted (**Fig. 1A**). Details for each test are found below.

**Figure 1.**
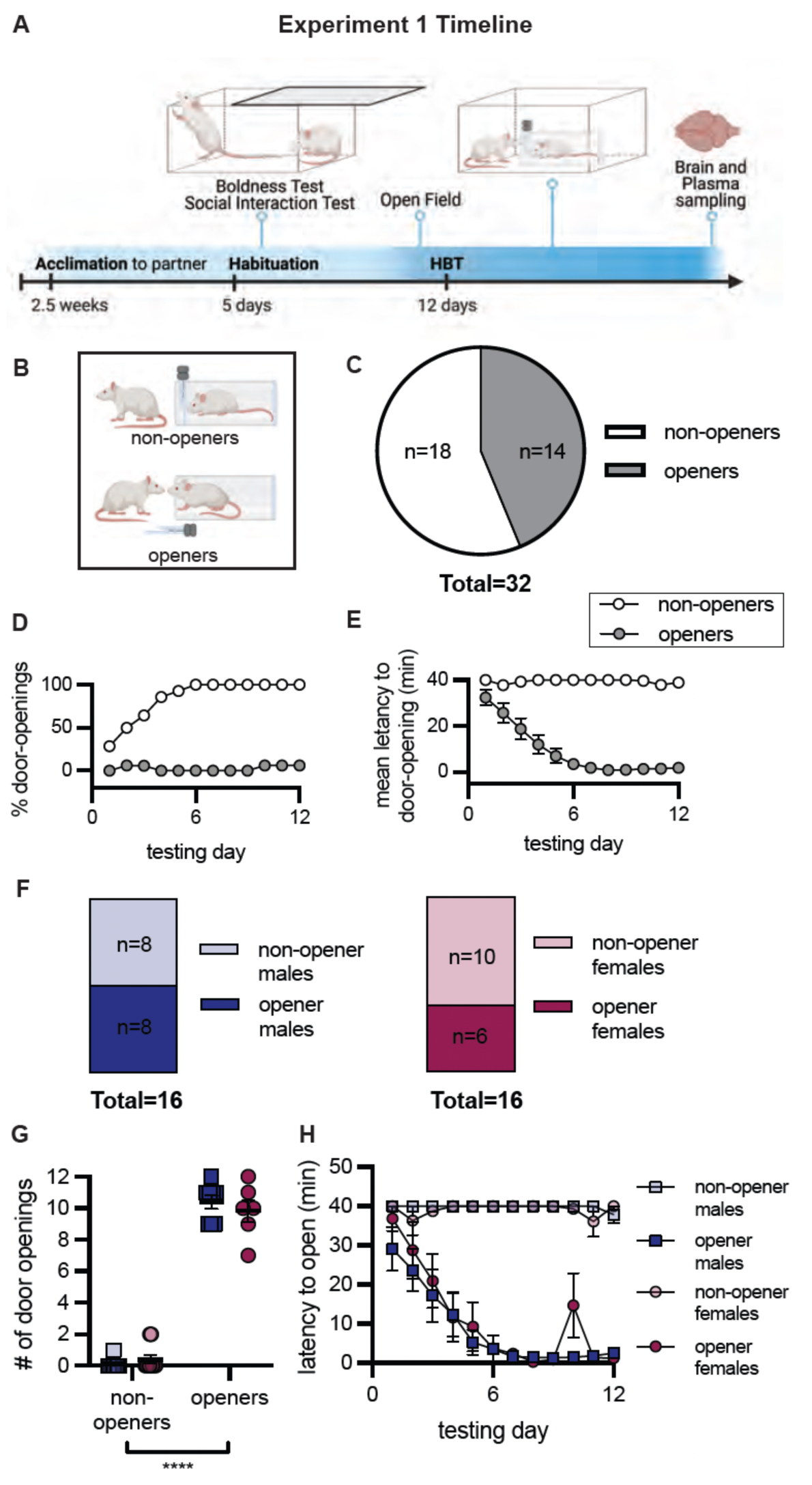
Adult helping behavior is similar in male and female rats. A. Experimental Timeline. During habituation, animals underwent a boldness test, a social interaction test and an open field assay. The helping behavior test (HBT) consisted of 12 days of 1-hour sessions. On the final session, brains and plasma were extracted for processing. B. Animals were categorized into ‘openers’ or ‘non-openers’ according to their behavior in the HBT. C. Percent of openers across all animals: 43.75% (14/32) of rats became openers. For openers, helping behavior consisted of increased % of door-openings (D) and decreased latency to open (E) across testing sessions. F. A similar percent of male rats (50%, 8/16) and female rats (37.5%, 6/16) became openers. The number of door openings across the 12 testing sessions did not differ by sex (G), nor did the latency to open (H). Data are mean ± SEM.

##### Boldness Test

The Boldness Test was used to reduce differences in door opening caused by the rats’ hierarchy within the pair, or by individual traits such as curiosity, and anxiety-like behavior (Ben-Ami Bartal et al., 2011). For five consecutive days, the metal grid top of the home cage was opened halfway, and the time it took for each rat to go to the open half, place its two front paws on the edge of the cage and peek out was recorded. The test ended after 5 minutes if both rats did not peek out. The rat who peeked first at least 3 of the 5 days was assigned the ‘free rat’ role and the other cagemate the ‘trapped rat’ role. This protocol was conducted according to (Ben-Ami Bartal et al., 2011). The mean latency to peek across the five days was calculated for each rat, as well as the difference, or delta, between the (future) free rat and the (future) trapped cagemate.

##### Social Interaction Scoring

Social interactions (SI) were measured prior to starting the HBT to characterize social affiliation of each pair (Panksepp and Beatty, 1980; Kondrakiewicz et al., 2019). Each cagemate pair was placed together in the testing arena for 30 minutes. In the first 5 minutes, several social interaction measures were recorded, including: the frequency and duration for each pair of sniffing, lying together, following, and climbing. Total social interaction time and number of social interactions were also measured on the first session of the HBT, during the 5 minutes after the door opened (whether by the free rat or by the trapped rat after the half-way door opening).

##### Open Field Test (OFT)

The OFT was used to analyze the rats’ relative approach-avoidance behavior (Walsh and Cummins, 1976; Saenz et al., 2006; Doron et al., 2012). Each rat entered the testing arena alone and underwent the OFT for 30 minutes. The following measures were taken: activity (measured by the number of cm the rat walked in the arena) and time in the center of the arena (the center was a quarter of the total arena size – 25 × 25 cm).

##### Helping Behavior Test (HBT)

For the HBT, a restrainer was added to the center of the testing arena (Harvard Apparatus, Holliston, MA, USA). The restrainer, made of transparent Plexiglas (9 w X 19.7 d X 8.26 h cm), had several slots so that communication between the rats was possible via sight, smell, hearing, and touch. The restrainer had a homemade door that could only be opened from the outside and, therefore, only by the free rat. There were two weights on the door, totaling 50 g. Because of the weights, a deliberate effort was required to open the door; a rat who wanted to open it and knew how to open it would succeed, however, the door would not open accidentally.

Following habituation, rats underwent 12 consecutive days of HBT testing (except Saturday; **Fig. 1A**). On each day, the free rat was placed into the arena once the restrainer with the trapped rat was set. The free rat had 40 minutes to open the door and release the trapped cagemate. Even though the door was designed to be opened exclusively from the outside, some trapped rats managed to open it from the inside (n=12, 37.5%). In this case, the trapped rat was returned immediately to the restrainer, and a blocker was added, preventing the trapped rat’s access to the door. The blocker was then also used on all forthcoming days of testing. If the free rat opened the door, the experimenter removed the blocker immediately. After 40 minutes, if the free rat did not open the door, the experimenter opened the door halfway (to a 45° angle) for another 20 minutes. If there was a blocker, it was removed at this point. The partial opening encouraged the free rat to open the door and allowed the trapped rat to open it from the inside, avoiding learning helplessness. Only door opening that occurred by the free rat in the first 40 minutes was considered as door opening for analysis. At the end of the experiment, the number of total door openings was calculated for each pair. Based on previous studies (Ben-Ami Bartal et al., 2011, 2021), pairs in which the door was opened at least twice on the last three days were classified as ‘openers’. After classification, the percentage of rats that opened the door on each testing day was calculated per group. In addition, the average time to door-opening was calculated (when the door was not opened, a 40-minute (maximum test time) score was given). On the first and last session, the following measures were recorded in the first 40 minutes: velocity, time in the corners, time around the restrainer, number of entries to the corners and number of entries into the restrainer area. These measures were recorded only for rats that did not open the door in order to allow for statistical comparison.

#### Biological Measures

##### Plasma and brain section collection

Three days after the paradigm ended, all rats were sacrificed by rapid decapitation following a brief CO2 exposure. Both brains and plasma were collected from the free rats, while only plasma was collected from the trapped rats. Brains were snap-frozen on dry ice, and stored at –80°C.

For plasma collection, trunk blood was first collected in EDTA tubes and centrifuged at 3000 rpm, under 4 ℃ for 15 minutes. Plasma was aliquoted and frozen at –80℃ until CORT analyses. According to the manufacturer’s protocol, plasma CORT concentrations were evaluated using an Enzyme-Linked Immunosorbent Assay (ELISA) kit (EC3001, AssayPro, St. Charles, MO).

Brains from 20 animals (10 openers, 10 non-openers – 5 of each sex) were sliced in coronal orientation on a cryostat. Tissue was obtained from each hemisphere using a 1mm diameter Miltex biopsy puncher (Bar Naor, Israel). One hemisphere’s brain tissue punch was utilized for RNA-sequencing, while the other hemisphere’s punches were preserved for real time quantitative PCR (rtPCR) analyses. Punches were taken directly while the brain was on the cryostat, and subsequent slices were used to confirm thickness of the punch. Brain sections of the NAc and AI were collected, according to the following coordinates (Paxinos and Watson, 2006: NAc: AP 2.2 to 1.2 mm; AI: AP 3.2 to 1.2). Tissue punches were stored in clean tubes and were immediately frozen on dry ice, and stored at –80°C.

### Gene expression profiling

#### RNA sequencing (RNA-seq)

Isolated NAc and AI punches were subjected to genome-wide transcriptional profiling at the at the Technion Genomics Center and the UCLA Social Genomics Core Laboratory, respectively.

For AI samples, RNA was extracted from 1mm diameter tissue punches of approximately 20 μg of frozen brain tissue (Qiagen RNeasy), assessed for suitable mass (RiboGreen), reverse transcribed to cDNA using a high efficiency mRNA-targeted enzyme system (Lexogen QuantSeq 3′ FWD) and subsequently sequenced using Illumina NovaSeq instrument (Lexogen Services, GmbH). Sequencing targeted at least 10 million sequencing reads per sample (achieved mean = 17.7 million), each of which was mapped to the mRatBN7.2 genome sequence (average 97.9% mapping rate) and normalized to transcripts per million using the STAR aligner.

For NAc samples, RNA was extracted from 1mm diameter tissue punches of approximately 20 μg of frozen brain tissue using the QIAcube Connect with an RNeasy micro kit (Qiagen) and assessed for suitable mass using an Agilent TapeStation System. The RINe values of all samples were in the range of 7.9-9.1, indicating a high quality. RNAseq libraries were constructed using a NEBNext Ultra II Directional RNA Library Prep Kit (New England Biolabs). mRNA pull-down was performed using the Magnetic Isolation Module (New England Biolabs). After construction, the concentration of each library was measured using a Qubit (Invitrogen) and the size was determined using the TapeStation 4200 with a High Sensitivity D1000 kit (Agilent). All libraries were mixed into a single tube with equal molarity. RNAseq data was generated using on an Illumina NextSeq instrument (Lexogen Services, GmbH) using P2 100 cycles (Read1-100; Index1-8; Index2-8). Quality control was assessed using Fastqc (v0.11.5), reads were trimmed for adapters, low quality 3’ and minimum length of 20 using CUTADAPT (v1.12). 100 bp single reads were aligned to a rat reference genome (Rattus_norvegicus.Rnor_6.0.faENSEMBL) and normalized to transcripts per million using STAR aligner.

#### Quantitative real-time polymerase chain reaction (rtPCR)

Total RNA was first isolated with Trizol (Thermo Scientific) and chloroform (Sigma-Aldrich Israel Ltd.). Subsequently, 25ng of RNA per reaction was reverse transcribed into cDNA using the High Capacity Reverse Transcription Kit with RNase Inhibitor (Thermo Scientific). The quantitative real-time polymerase chain reaction (PCR) was conducted using a Fast SYBR Green PCR Master Mix (Applied Biosystems) along with specific primers for *Oxtr* and GAPDH genes (HyLabs Israel Ltd.). To standardize gene expression levels, *Oxtr* expression was normalized to GAPDH, as the housekeeping reference gene. Product purity was validated through a melt curve analysis using ABI hardware and software (QuantStudio Real-Time PCR Systems, Thermo Scientific), and gene expression analyses were determined using the comparative ΔΔCt (fold change) method.

#### Immunohistochemistry

To examine neuronal activity associated with helping behavior, c-Fos expression was analyzed in a separate group of rats (Experiment 2). In brief, all animals performed the HBT experiment in similar conditions to the Wistar rats in Experiment 1, described above. However, several weeks before the habituation section, rats received a stereotactic injection of the retrograde tracer, Fluoro-Gold into the NAc (see Ben-Ami Bartal et al., 2021 for complete methods), and were allowed to recover before starting the behavioral tests. Moreover, on the last HBT session, the restrainer door was locked, and rats were perfused immediately after the session ended. An average of 24 slices per animal (range 15-28 slices) from atlas coordinates AP +5.2 to –8.3 (Paxinos & Watson, 2006) were analyzed with the automated software Brainways developed in house (Kantor and Bartal, 2023). The Brainways software allows automatic analysis of histological brain slices. Using this pipeline, slices were first matched to the atlas coordinates. Minor manual fine-tuning was then done by an experimenter blind to experimental conditions to optimize accuracy. At the next stage, the software automatically ran a cell detection algorithm, detecting c-Fos expression over the different slices. Then, cell density (number of c-Fos+ cells per 250^2^ μm) was calculated for each brain area. Missing values were interpolated by the average value of all samples, for each condition.

### Statistical Analyses

#### RNA-seq analysis

The differential expression of each gene (DEG) was estimated with a standard linear statistical model relating log2-transformed transcript abundance values to measure individual status as opener vs. non-opener while controlling for sex. *A priori* hypotheses were generated for key genes related to social behavior and reward, including for genes related to OXT, dopamine (DA) and CRH receptors, as well as genes associated with early immediate genes like *Fos*. A full list of these genes can be found in **Supplemental Table S1**. Fold change analyses comparing openers and non-openers was conducted on each of these genes.

We next applied a bioinformatic analysis of transcription factor-binding motifs (TFBM) in core promoter sequences of the DEGs, using the Transcription Element Listening System (TELiS, http://www.telis.ucla.edu/) on all genes showing ≥ 1.5-fold differential expression in opener vs. non-opener animals. *A priori* hypotheses were generated for key transcriptional regulators related to immediate early genes and stress pathways, such as KROX, API, CREB, and SP1. For all bioinformatics analyses, standard errors were computed and estimated by bootstrap resampling of linear model residual vectors across genes (200 cycles); this controls for any statistical dependence amongst genes (Cole et al., 2005; Powell et al., 2013).

#### Task PLS analysis

The multivariate task partial least square (PLS) analysis was based on brain regions’ c-Fos expression to identify optimal neural activity patterns that distinguished between the experimental groups (McIntosh et al., 1996; Mcintosh, 1999). Task PLS looks for latent variables (LVs) that explain a significant portion of the data variability. Through singular value decomposition, PLS produces a set of mutually orthogonal LV pairs. One element of the LV depicts the contrast, which reflects a commonality or difference between conditions. The other element of the LV, the relative contribution of each brain region (termed here ‘salience’), identifies brain regions that show the activation profile across tasks, indicating which brain areas are maximally expressed in a particular LV. Statistical assessment of PLS was performed by using permutation testing for LVs and bootstrap estimation of standard error for the brain region saliences. For the LV, significance was assessed by permutation testing: resampling without replacement by shuffling the test condition. Following each resampling, the PLS was recalculated. This was done 500 times in order to determine whether the effects represented in a given LV were significantly different than random noise. For brain region salience, reliability was assessed using bootstrap estimation of standard error. Bootstrap tests were performed by resampling 500 times with replacement, while keeping the subjects assigned to their conditions. This reflects the reliability of the contribution of that brain region to the LV. Brain regions with a bootstrap ratio >2.57 (roughly corresponding to a confidence interval of 99%) were considered as reliably contributing to the pattern. Missing values were interpolated by the average for the test condition. An advantage to using this approach over univariate methods is that no corrections for multiple comparisons are necessary because the brain region saliences are calculated on all of the brain regions in a single mathematical step. MATLAB code for running the Task PLS analysis is available for download from the McIntosh lab website.

#### Network analysis

Network maps were created using a correlation matrix of c-Fos+ cells between all brain regions (using pairwise Pearson correlation coefficients calculations). Based on scale-free network characteristics described previously (Ben-Ami Bartal et al., 2021; Breton et al., 2022), only the top 10% correlations were used to produce the network graphs. Correlation values higher than the cutoff were set to one and the corresponding brain regions greater than 1 were considered connected to the network. For more detailed methods see (Ben-Ami Bartal et al., 2021).

#### Additional statistical analyses & details

The variables ‘group’ (openers, non-openers) and ‘sex’ (male, female) were analyzed as between-subject variables, and the variables ‘role’ (free, trapped) and ‘days of test’ (1 to 12) were analyzed as within-subject variables. A chi-square test of independence was used to analyze if there was a difference in the number of rats that became openers based on sex or experimental batch. A Friedman’s test was used to test for differences in the proportion of door-opening across groups. All other measures were analyzed with analysis of variance (ANOVA) or t-tests, as appropriate. Sidak’s post hoc tests were used for multiple comparisons. All tests were two-tailed with a significance level set at p<0.05. Results are displayed and reported as means ± the standard error of the mean (SEM). Statistical analyses were conducted using SPSS (version 26), Prism (version 9; Graphpad Software), or MATLAB with α set to 0.05.

## Results

### Adult male and female rats behaved similarly towards trapped cagemates during the HBT

In Experiment 1, 32 pairs of adult male and female Wistar rats were tested in the HBT with trapped cagemates of the same strain and sex over a two-week period (**Fig. 1A**). During the 1-hour testing sessions, free rats could open the restrainer door thereby releasing their trapped cagemate. Based on their door-opening behavior, free rats tested in the HBT were classified as ‘openers’ if they learned to release the trapped rat and demonstrated persistent door-opening over consecutive testing sessions, or ‘non-openers’ if they failed to do so (**Fig. 1B**; see methods). Out of all pairs tested, 43.75% rats were classified as openers (n=14/32, mean door-opening=10.21±0.36) and 56.25% rats were classified as ‘non-openers’ (n=18/32, mean door-opening=0.28±0.16) (**Fig. 1C**; see **Fig. S1A** for individual door opening data). For openers, the proportion of door-openings significantly increased (Friedman’s test: p<0.0001, **Fig. 1D**) and latency to door-opening decreased (F(2.5, 32)=21.30, p<0.0001, **Fig. 1E**) along the testing days, which was not the case for non-openers (p>0.05). A similar proportion of openers was observed for males and females, with 8/16 of males and 6/16 of females becoming openers (Fisher’s exact test: p=0.7224, **Fig. 1F**). A 2-way ANOVA examining the effects of sex and opener status on the number of door-openings identified a main effect of opener status (F(1,28)=656.5, p<0.0001) but no main effect of sex and no interaction between sex and opener status (p>0.05, **Fig. 1G**). Similarly, a mixed-effects model testing the effects of sex and opener status on door-opening latency across the testing days identified a main effect of time (F(11,297)=18.36, p<0.001), opener status (F(1,28)=474.3, p<0.0001 and a significant interaction between time and opener status (F(11,297)=18.96, p<0.0001), but no effect of sex and no interactions between sex and opener status (**Fig. 1H**). As there were no sex effects in these primary analyses, males and females were grouped together for all subsequent analyses. In sum, about half of the animals learned to open the restrainer by the end of testing, with no observed sex differences.

### Opener rats showed greater affiliation with their trapped cagemate

We next assessed if behaviors prior to the start of the HBT could predict who would subsequently become openers. In a manner consistent with the criteria for the division into the free and trapped roles, in a 2-way ANOVA, there was a main effect of rat role whereby in the Boldness Test, the (future) free rats peeked faster than the (future) trapped rats (F(1,28)=24.567, p<0.0001; **Fig. 2A-B**). More importantly, there was also a main effect of opener status, whereby future openers (free and trapped rats) peeked faster than future non-openers (F(1,28)=4.964, p=0.034; **Fig. 2B**). There was also a trend for an interaction between rat role and subsequent opening status (F(1,28)=3.996, p=0.055), though this did not reach statistical significance. In a post-hoc test, among the (future) trapped rats, rats from the openers group peeked faster than those in the non-openers group (p=0.026; **Fig. 2B**). However, among the (future) free rats, there was no significant difference between openers and non-openers (p=0.216; **Fig. 2B**). We next calculated the difference in peeking latency within each pair of cagemates (**Fig. 2C**). Here, there was a statistically significant difference between the future non-opener and opener groups (t(23)=1.287, p=0.0389), with non-openers showing a greater discrepancy in peeking time between the two cagemates (27.4 ± 6.3 sec) relative to openers (11.9 ± 3.3 sec; **Fig. 2C**). There were no group differences in either velocity (**Fig. S1B)** or time spent in center (**Fig. S1C)** during open field testing, indicating that baseline movement and anxiety-like behavior are not likely explanations for these findings, nor do they predict subsequent opening.

**Figure 2.**
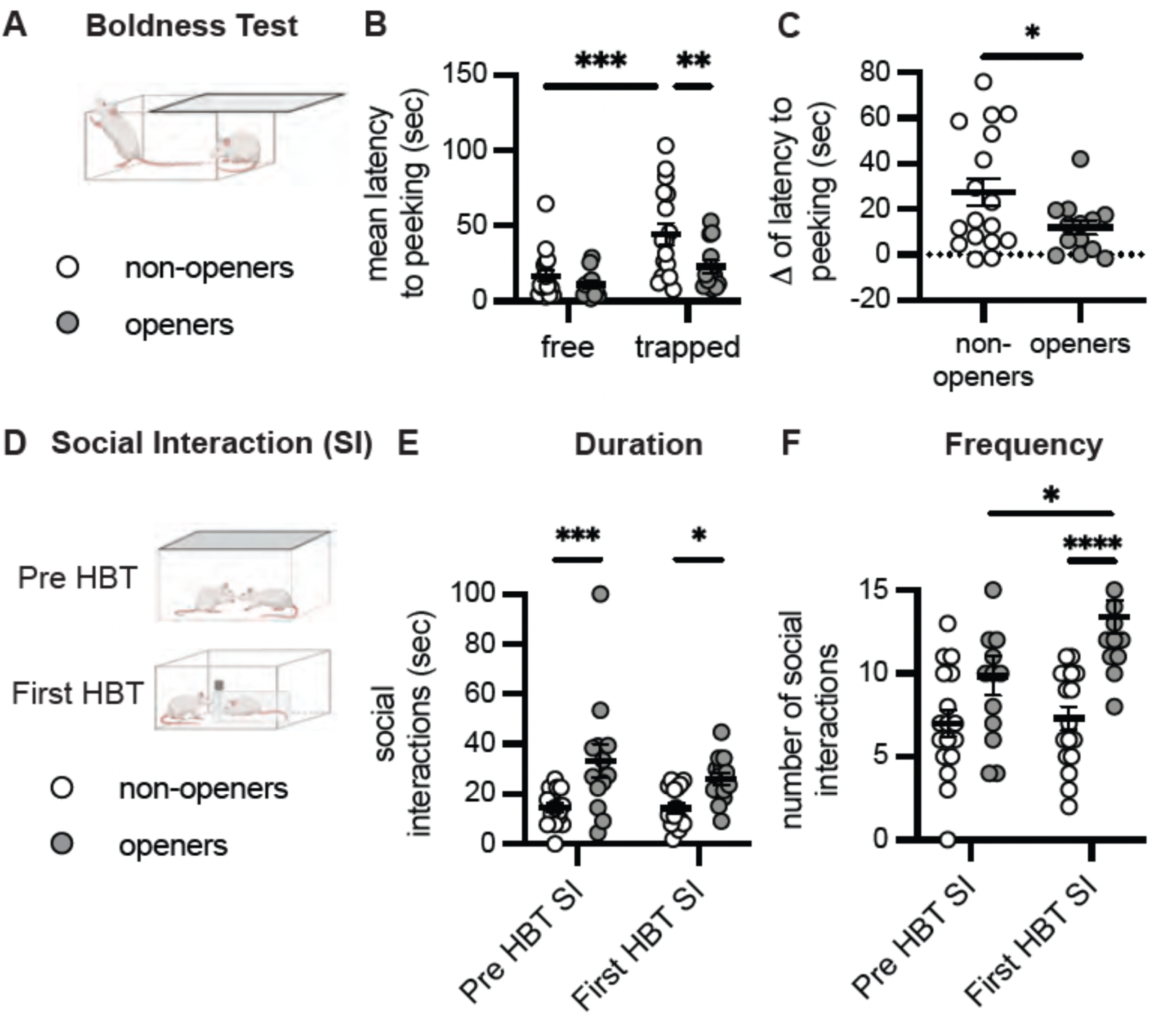
Social affiliation and boldness predict subsequent opening behavior. A. Diagram of the boldness test conducted prior to the HBT. B. Rats designated to become the ‘free’ rat had a faster latency to peak during the boldness test than future ‘trapped’ rats. C. ‘Opener’ pairs showed less of a difference in peeking latency within the cage than did ‘non-opener’ pairs D. Social interaction was scored during habituation, prior to the HBT (Pre-HBT), and on the first day of the HBT itself (First HBT), following door opening. Openers showed increased duration (E) and frequency (F) of social interactions, both prior to the HBT and on the first day of HBT testing. Data are mean ± SEM. * p<0.05. **p<0.01, ***p<0.001, ****p<0.0001.

Next, we looked at SI, both prior to the HBT, and on the first day of the HBT after the restrainer door had been opened (either by the free rat (n=4/32) or at the halfway point by the trapped rat (n=28/32 rats; **Fig. 2D**). A 2-way ANOVA comparing opener and non-opener groups at these two timepoints revealed a main effect of opener status on SI duration (F(1,54)=18.4, p<0.001 with no effect of session (Pre-HBT or Day 1 of HBT) and no interaction between them (**Fig. 2E**). Planned post-hoc tests indicated that overall, compared to the non-opener pairs, the pairs from the openers group spent a greater amount of time interacting with one another, both prior to the HBT and on Day 1 of the HBT (p=0.001 and p=0.044 respectively). This indicates that rats that would subsequently become openers displayed more affiliative social interactions at baseline. A robust main effect of opener status was also found for SI frequency (F(1,56)=24.29, p<0.001), as well as a main effect of session (F(1,56)=4.467, p=0.039) and a trend towards an interaction between them (F(1,56)=3.201, p=0.079; **Fig. 2F**). Planned post hoc tests indicated that openers had significantly more frequent social interactions than non-openers on Day 1 of the HBT (p<0.001); this effect was trending prior to the HBT (p=0.06) but did not reach statistical significance. Furthermore, there was a significant difference in the frequency of interactions across sessions, with openers, but not non-openers, showing an increased number of interactions on the first day of the HBT compared to the pre-test session (p=0.0241; **Fig. 2F**).

To further identify differences in motivational state between openers and non-openers, movement patterns prior to door opening were also analyzed on the first HBT session. This analysis revealed that openers and non-openers showed similar activity patterns around the trapped rat, including velocity, time spent in the arena corners, time around the restrainer, and number of entries to these regions (p>0.05 for all measures; **Fig. S1D**). This indicates a similar motivational state for rats tested with a trapped cagemate regardless of their subsequent pattern of helping behavior. Furthermore, while it is impossible to determine with certainty whether non-openers failed to open the restrainer from a lack of motivation or a lack of ability to learn the task, a significant reduction in their efforts and other-focused behavior was observed by the end of testing (day 12 – day 1; **Fig. S1E**) suggesting that non-openers were less perseverant.

Thus, affiliative behavior, both prior to the HBT, and that expressed towards the released rat on the first session, was a better predictor of subsequent door-opening than movement patterns around the trapped rat on the same session. Overall, this indicates that social interaction is a strong predictor of future opening behavior.

### Oxytocin receptor mRNA levels were elevated in the nucleus accumbens (NAc) of openers compared to non-openers

To identify genome-wide transcriptomic differences that correspond with opening behavior in the HBT, we utilized RNA-sequencing to measure the effect of opening behavior on gene expression in the NAc and AI; we focused on these two regions given their role in empathy (Wu and Hong, 2022), helping behavior (Ben-Ami Bartal et al., 2021) as well as social reward (Dölen et al., 2013; Dölen and Malenka, 2014) **(Fig 3A)**. Based on *a priori* hypotheses, we first examined changes in genes related to oxytocinergic (*Oxtr)* and dopaminergic (*Drd1, Drd2)* signaling, as well as genes related to the stress axis (*Crhr1, Nr3c1*) and immediate early gene activity (*Fos1l*) (**Fig. 3A, Table S1**). In particular, given the known role of oxytocin in social reward (Dölen et al., 2013; Dölen and Malenka, 2014), we hypothesized that oxytocin receptor gene expression would be differentially expressed in openers and non-openers. *Oxtr* expression was significantly upregulated in openers compared to non-openers in the NAc (2.6-fold change, p=0.006) but not in the AI (0.948-fold change, p=0.463). No significant differences in DA receptor or CRH receptor gene expression were observed in either the NAc or AI of openers compared to non-openers. However, a significant upregulation in *Nr3c1*, the gene encoding the glucocorticoid receptor, was found in the AI, but not in the NAc, of openers (1.19-fold change, p = 0.0289). Additionally, a marker for neuronal activation, the *Fosl1* gene, was significantly upregulated only in the NAc of openers (2.067-fold change, p = 0.0233; **Fig. 3A**). Together, this suggests that differences in OXT, but not DA or CRH, signaling in the NAc play a role in helping behavior.

**Figure 3.**
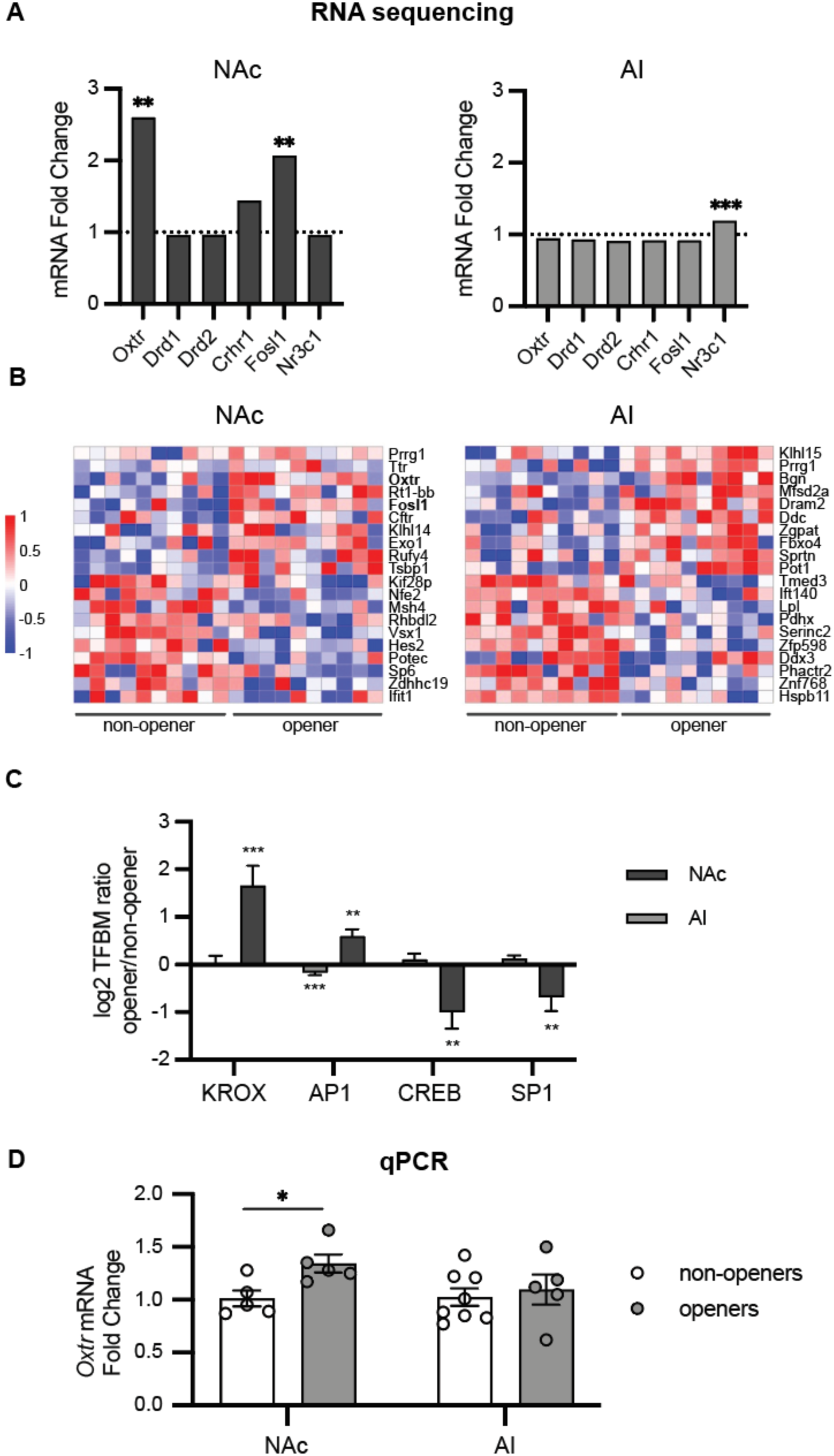
RNA analyses comparing openers and non-openers. A. RNA-sequencing of a-priori defined genes of interest in the NAc and AI. Increased oxytocin receptor and Fos gene expression levels were observed in the NAc of openers relative to non-openers. Data is shown as a fold change, with 1 indicating no difference between openers and non-openers and changes greater than 1 indicating elevated levels in openers relative to non-openers. B. Top 10 up-regulated and top 10 down-regulated differentially expressed genes (DEGs) in the NAc and AI. Red indicates up-regulation in openers relative to non-openers, while blue indicates down-regulation relative to non-openers. C. Transcription factor binding motifs (TFBM) prevalence in openers vs. non-openers. Positive numbers indicate increased prevalence in openers, while negative numbers indicate increased prevalence in non-openers. D. Real time quantitiative PCR (qPCR) results of oxytocin receptor mRNA levels in the NAc and AI. Data are presented as a fold change with non-openers having a mean of 1. * p<0.05, **p<0.01, ***p<0.001.

RNA-sequencing also allowed us to explore the relationship between opening behavior and transcriptome-wide gene regulation in these two regions. We conducted an analysis of differential gene expression (DEG) in NAc and AI samples from opener vs. non-opener animals while controlling for sex. This analysis identified 463 ≥ 1.5-fold DEGs in the NAc (226 up-regulated and 237 down-regulated) and 956 in the AI (593 up-regulated and 363 down-regulated). The top 10 up-regulated and down-regulated DEGs are shown in **Fig. 3B**. Importantly, in the NAc, both *Oxtr* and *Fosl1* were in the top 10 up-regulated genes found in openers. Of note, *Prrg1*, a gene encoding a vitamin-K dependent transmembrane protein, was the only gene significantly up-regulated in both the AI and NAc of openers. Though the function of many of these genes are still being explored, together, these genes provide candidate targets that may contribute towards opening behavior.

Next, we applied a bioinformatic analysis of transcription factor binding motifs (TFBMs) in core promoter sequences of genes differentially expressed in the NAc or AI of opener vs. non-opener rats (**Fig. 3C**) using the Transcription Element Listening System (TELiS) (Cole et al., 2005). Two transcription factors, KROX and AP1, were looked at *a priori* given their association with immediate-early gene (IEG) activity (Dragunow, 1996; Kovács, 2008). This analysis showed a significant increase in KROX and AP1 activity in openers in the NAc (1.65±0.418, p=0.001 and 0.599±0.139, p=0.0001, respectively; **Fig. 3C**). However, in the AI, there was no significant change in KROX activity (0.028±0.153, p=0.85), while AP1 activity showed a small but significant decrease (−0.169±0.049, p=0.001; **Fig. 3C**). As both KROX and AP1 transcription factors are associated with IEG activity, the simultaneous elevation of these factors in the NAc suggests a higher level of neuronal activation in the NAc in rats with a history of opening. Furthermore, these findings align with the observed upregulation of the *Fosl1* gene in the NAc (**Fig. 3A**). In addition, both KROX and AP1 appeared in the top 25 up-regulated TFBMs in the NAc (for a full table, see **Supplemental Table S2**). CREB was another key transcription factor of interest, given the CREB/ATF family’s broader role in emotional regulation (Green et al., 2008) and prior work observing stress-associated upregulation of CREB in the NAc (Barrot et al., 2002; Muschamp and Carlezon, 2013). In the NAc, there was significantly decreased CREB activity for openers compared to non-openers (−0.996±0.352, p=0.01), with no significant difference in the AI (0.108±0.124, p=0.38; **Fig. 3C**). Further, ATF family transcription factors were among the top downregulated TFBMs in the NAc (**Table S2**). Notably, SP1, a transcription factor associated with oxidative stress (Ryu et al., 2003) was significantly decreased in the NAc of openers (– 0.687±0.290, p=0.02), with no significant difference in the AI (0.121±0.0710, p=0.09; **Fig. 3C**).

To validate our RNA-seq findings, we next performed qPCR to analyze *Oxtr* mRNA levels in the AI and NAc, using tissue from the same animals as the RNA-seq study. (**Fig. 3D**). In line with the RNA-seq results, qPCR revealed a significant increase in *Oxtr* expression in the NAc (t(8)=2.91, p=0.02) but not the AI (p>0.05) of openers compared to non-openers (**Fig 3A)**. In sum, across two different methods, we found substantial gene expression changes in the NAc, with increased *Oxtr* expression in animals with a history of opening behavior; this aligns with our previous knowledge of the NAc and its known role in helping behavior (Ben-Ami Bartal et al., 2021).

### Increased c-Fos was observed in social neural networks of openers compared to non-openers

In order to map the neural circuits activated during the HBT in openers and non-openers, the immediate-early gene c-Fos was analyzed across the brain in a separate group of adult SD male rats (n=13) tested with a trapped SD cagemate for two weeks (Experiment 2, **Fig. 4A**). On the final testing session, the restrainer was latched shut, and c-Fos^+^ cells were measured as an index of the neural activity associated with an hour of being in the presence of a trapped cagemate (**Fig. 4B**). This strategy has previously been useful for us and others for describing neural networks involved in complex activity (Wheeler et al., 2013; Vetere et al., 2017; Ben-Ami Bartal et al., 2021; Breton et al., 2022). In this group, similar to Experiment 1, most rats (n=8/13) exhibited robust door-opening for trapped cagemates (average learning day: 5.13±0.9) and were classified as “openers”. Additionally, one rat opened on 2 consecutive days, as well as on the final testing day and was also categorized as an “opener”; thus, there were 9 openers in total (n=9/13, 69.3%). Several rats (n=4/13, 30.7%) rarely opened the restrainer and did not show consecutive door-opening behavior; these were classified as “non-openers” (**Fig. 4C-E**).

**Figure 4.**
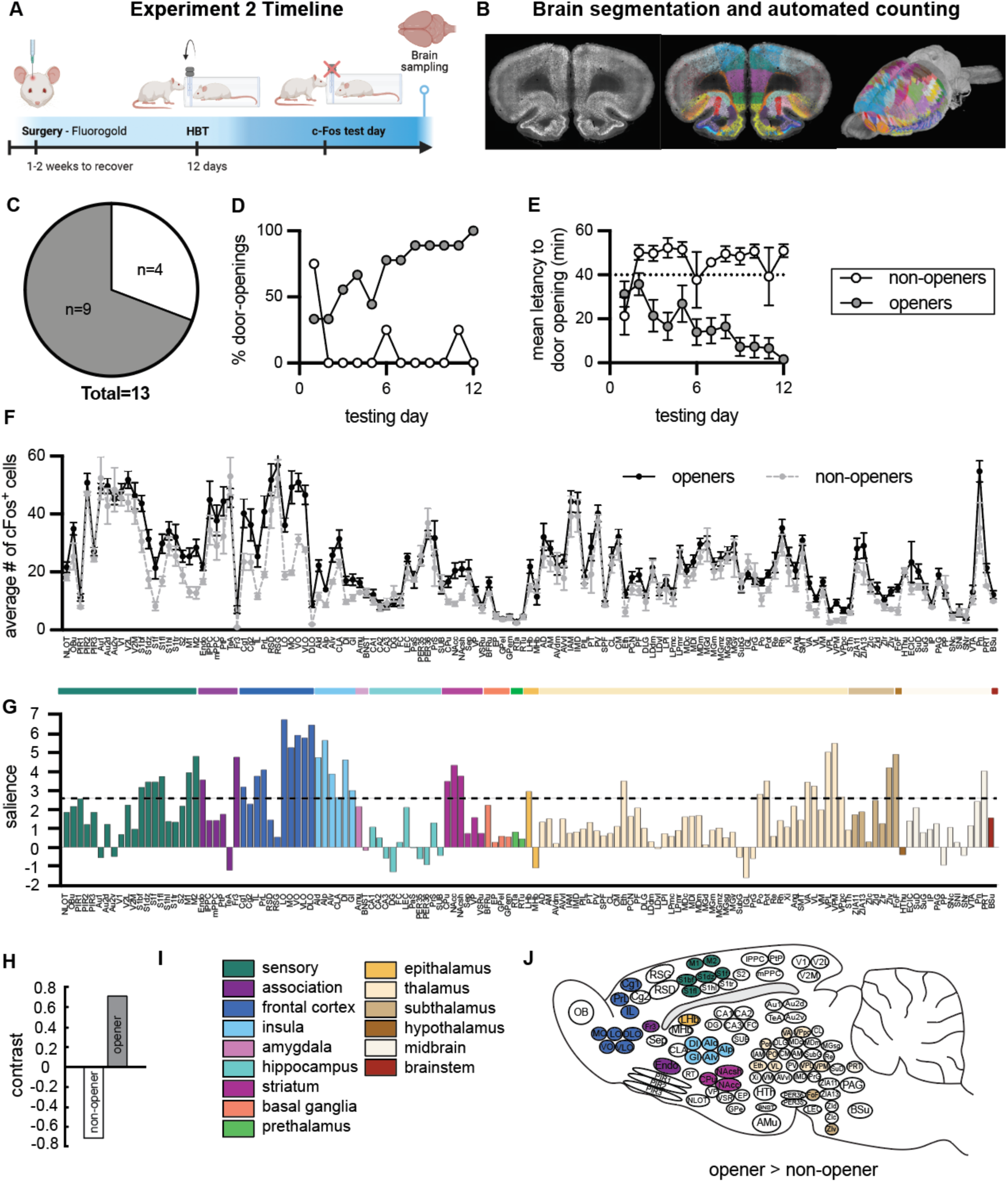
Brain-wide patterns of neural activity in openers. A. Experimental Timeline. These same animals were used in analyses found in (Bartal et al., 2021). B. Representation of c-Fos analysis pipeline, following (Kantor and Bartal, 2023). C. Percent of rats that became openers (69.3%, 9/13) was similar to Experiment 1. Percent-door openings increased (D) and latency to open decreased (E) across testing days for openers. F. Number of c-Fos+ cells per 250 μm region (mean±SEM) for opener and non-opener rats. G. Partial least square (PLS) task analysis. A history of opening in the HBT was associated with increased activity in multiple brain regions compared to all other conditions. Regions that cross the dashed significantly (p<0.05) contributed to this pattern H. Openers and non-openers showed distinct patterns of neural activity (PLS contrast) I. Legend of brain region categories coded by color. J. Diagram of rat brains showing regions significantly more activity in openers relative to non-openers.

Using an in-house software (Kantor and Bartal, 2023), c-Fos^+^ was quantified in 24 slices per rat on average, and the number of c-Fos+ cells was compared between openers and non-openers (**Fig. 4B, F**, see methods for details). In total, 137 regions were analyzed (**Table S3**). In order to compare activity levels across regions, c-Fos^+^ cell numbers were normalized to a standard area of 250µm^2^. Analysis of brain-wide c-Fos^+^ patterns using a multivariate task Partial Least Square (PLS) approach (**Fig. 4G-J**) (McIntosh et al., 1996; Mcintosh, 1999) found a significant latent variable (LV) for the contrast between c-Fos expression across the brain of openers and non-openers (LV, p<0.05; **Fig. 4H**). This reflects higher brain-wide c-Fos^+^ cell numbers in the openers compared to the non-openers (t-test, t=7.26, p<0.0001, **Fig. 4F**). Bootstrapping and permutation tests were used to discover the pattern of neural activity associated with this contrast (**Fig. 4G**, see methods). Multiple brain regions significantly contributed to the contrast between openers and non-openers, including primary and secondary sensory regions such as somatosensory, motor, and olfactory cortices (**Fig. 4G**, in green). Additionally, the orbitofrontal regions, anterior cingulate cortex, mediofrontal regions (infralimbic IL, prelimbic PrL), insula, claustrum, lateral habenula (Lhab), ventral and posterior thalamic nuclei, subthalamic regions and midbrain regions all contributed to this contrast (**Fig. 4G**). This analysis indicates that in the presence of a trapped cagemate, openers demonstrated significantly increased activity in a dispersed network of brain regions that has previously been associated with empathy and prosocial motivation (Wu and Hong, 2022), and provides further support for the idea that this network supports a prosocial response towards conspecifics in distress. In addition, this analysis expands on previous work to include new brain regions and indicates a role for the IL, zona incerta (ZI), as well as specific thalamic areas such as the posterior nucleus (PO), in prosocial motivation **(Fig. 4J).** Interestingly, as opposed to previous findings comparing ingroup and outgroup conditions (Ben-Ami Bartal et al., 2021), here we did observe significantly increased activation in the anterior insula (AI) and anterior cingulate cortex (ACC) of openers compared to non-openers (**Fig. 4J**). Finally, c-Fos in the orbitofrontal cortex (OFC) contributed most strongly to the contrast between openers and non-openers, suggesting that this region plays a key role in predicting helping behavior.

### The ACC is a key node of openers’ neural functional connectivity network

In order to identify the functional connectivity involved specifically in adult helping behavior, a network analysis was conducted based on correlations between c-Fos^+^ cells in all brain regions of helper (opener) rats. Using Pearson’s pairwise correlation matrix and Louvain algorithm for clusterization (Breton et al., 2022), network graphs were generated based on the top 10% correlations (**Fig. 5A-B**, see methods for details). Connecting lines in the network graph represent high interregional correlations. The inter-region correlation matrix resulting from this analysis revealed four main clusters (**Fig. 5A**). Cluster 1 (**Fig. 5A**, blue) was composed mainly of sensory regions (auditory, visual, motor, & somatosensory cortices), as well as the insular cortex, some hippocampal regions (DG, CA3, perirhinal), the medial geniculate and the caudate and putamen (CPu). Cluster 2 (**Fig. 5A**, yellow) was composed of core regions of the prosocial response network described above (**Fig. 4**) as well as previously (Ben-Ami Bartal et al., 2021; Breton et al., 2022), including the NAc shell and core, OFC (VO, MO, VLO, LO, DLO), AI (Aid, AIv), PrL, claustrum, amygdala, lateral and medial habenula, ventral pallidum, olfactory regions, basal forebrain (an area that includes the band of Broca), and the frontal association area. Notably, here, the frontal association area was functionally connected to the amygdala and insula, as was previously reported (Nakayama et al., 2015). Cluster 3 (**Fig. 5A**, green) included mainly thalamic regions, the periaqueductal grey (PAG) region, and regions of the zona incerta (ZI). Lastly, Cluster 4 (**Fig. 5A**, pink) included frontal cortex regions such as the ACC and IL, the retrosplenial cortex (RSG), some hippocampal regions (CA1, CA2, SUB), the septum, BNST, thalamic & subthalamic nuclei, hypothalamic regions, as well as regions of the midbrain (substantia nigra (SN) & ventral tegmental area (VTA)) and brainstem. This cluster was most centrally located on the network and served as a connection point between the three other clusters. In particular, one region of Cluster 4, the ACC, has received a lot of attention for its role in empathy for distress, both in humans (Decety et al., 2016) and rodents (Jeon et al., 2010; Carrillo et al., 2019; Smith et al., 2021). A specific focus on the connectivity of just the ACC (**Fig. S2**) showed high correlations between the ACC and regions within all of the primary clusters, placing it as a hub within the network. Notably, the ACC was strongly connected to core regions of the prosocial behavior network, including the NAc shell and core, septum (Sep) prefrontal regions (IL, PRL), as well as the VTA and PAG. The projection between the ACC and NAc in particular has previously been reported to play a role in empathy for pain in mice (Smith et al., 2021) and prosocial behavior in the HBT (Ben-Ami Bartal et al., 2021), and our findings here bolster support for the role of this pathway in helping behavior. In sum, these findings support the validity of using c-Fos correlations for examining functional networks involved in the HBT, and identify a specific network involved in opening behavior.

**Figure 5.**
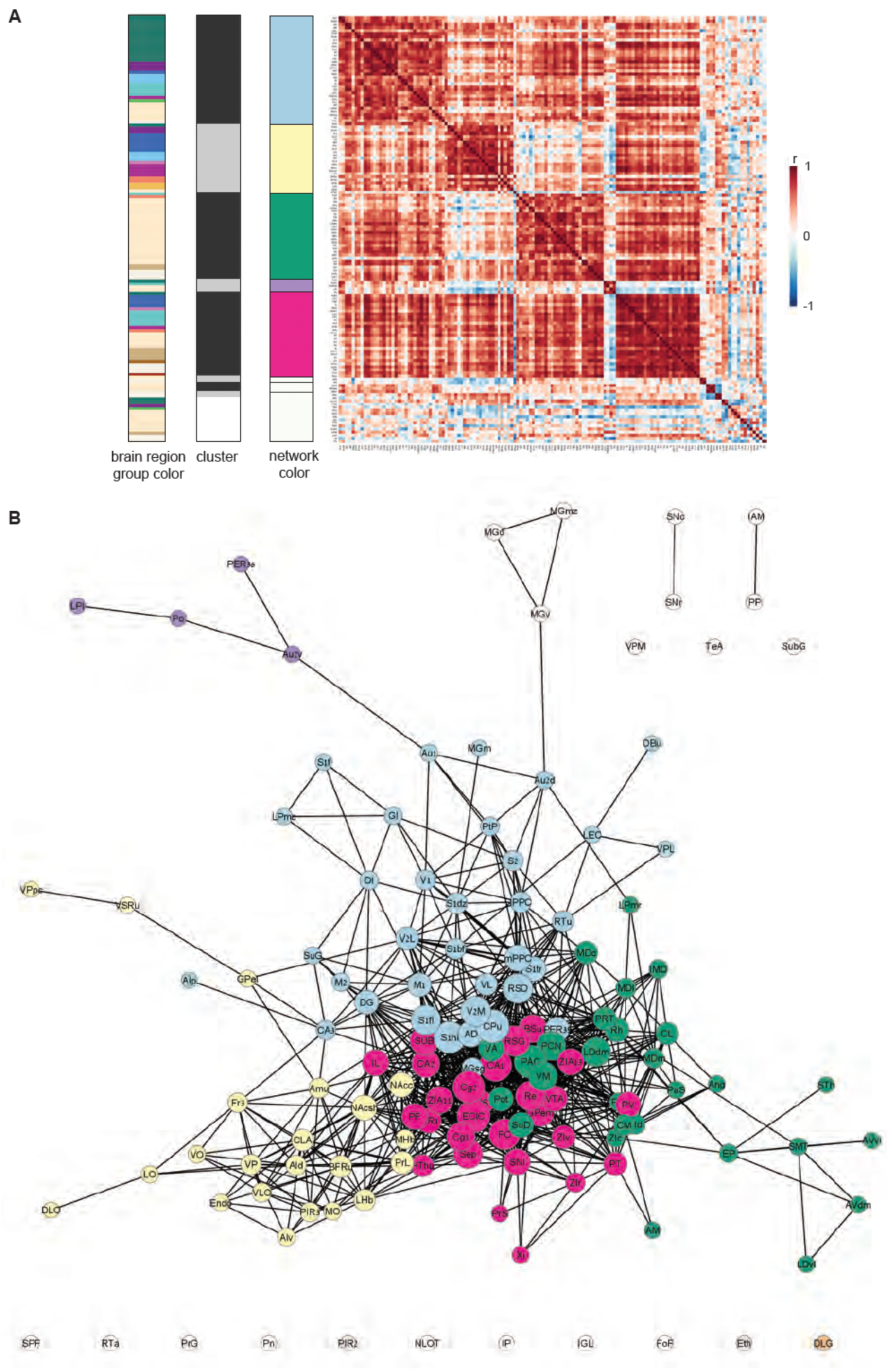
Network analysis of openers. A. Correlation matrix of c-Fos levels across all analyzed brain regions in openers. Red indicates a positive correlation, blue indicates a negative correlation. Four major clusters were identified. Coloring according to brain region group (see Figure 3I for legend), and according to the network map, is shown. B. Network map of openers. Solid lines connecting regions represent the top 10% of positive correlations.

## Discussion

Prosocial helping is observed across a wide range of species; however, even within the same social context, helping does not always manifest. These experiments took advantage of variability within a rodent model of helping behavior to explore behavioral and molecular changes that correspond with helping. Adult Wistar rats were tested with conspecifics of the same strain, a condition that typically elicits prosocial motivation (Ben-Ami Bartal et al., 2014). In this cohort, almost half of the animals demonstrated door-opening, with no differences across sexes. Social interactions measured both prior to and on the first day of restrainer testing differentiated openers from non-openers, with greater affiliative behaviors observed in animals that ultimately learned to open the restrainer. *Oxtr* expression, hypothesized to play a role in these differences, was quantified in the NAc and AI, two key regions of the prosocial brain network (Ben-Ami Bartal et al., 2021; Breton et al., 2022). Gene expression analyses identified elevated *Oxtr* expression in the NAc, but not AI, of openers, and elevated glucocorticoid receptor gene expression levels in the AI, but not the NAc. Brain-wide measurements of c-Fos in a cohort of SD rats indicated heightened neural activity for openers relative to non-openers in the prosocial brain network, including more activation in NAc, prefrontal, insular and sensory regions, with high levels of connectivity across these nodes. In sum, these findings indicate that affiliative behavior in dyads was associated with probability of helping and activation in the prosocial brain network.

Less helping was observed in dyads that exhibited fewer social interactions. This finding is in line with human literature; prosocial motivation is influenced by affiliation (Wilbanks et al., 2005) and prosocial behavior is more likely to be extended to closely affiliated others (Batson, 2011). While affiliative behavior has been demonstrated to influence helping in primates (Cronin, 2012), to our knowledge, the current study is the first to examine this in rats.

The association of affiliation with helping may indicate that, in highly affiliated pairs, increased social reward experienced from post-release contact motivates door-opening. Yet previous studies have demonstrated that social contact is not required for helping to occur (Ben-Ami Bartal et al., 2011; Sato et al., 2015; Cox and Reichel, 2020). Thus, although social contact may play a role in motivating helping, it is unlikely to completely explain this behavior. Alternatively, rats in affiliated pairs may find the conspecifics’ distress more salient or place a higher value on alleviating their distress, an effect that could be mediated by OXT signaling. For instance, pup retrieval has been shown to depend on an OXT-driven increase in synchronization in auditory cortex (Marlin et al., 2015; Carcea et al., 2021). Additionally, in a recent study, OXT receptor antagonism in the ACC delayed, but did not fully suppress, helping behavior in a variation of the HBT (Yamagishi et al., 2020).

Here, *Oxtr* expression was elevated in the NAc of opener rats relative to non-openers, in line with evidence for the role of NAc *Oxtr* in social approach (Yu et al., 2016; Williams et al., 2020), affiliative behavior (Ross and Young, 2009; Burkett et al., 2016; King et al., 2016) and social reward (Dölen et al., 2013; Hung et al., 2017). No difference was observed in the AI, mirroring prior work showing *Oxtr* expression in the NAc, but not AI, predicts social attachment (King et al., 2016). However, AI OXT blockade has been shown to reduce approach to a distressed conspecific (Rogers-Carter et al., 2018). *Drd1* and *Drd2* genes were also not different between openers and non-openers in these regions, despite their reported role in social play (Manduca et al., 2016), social attachment (Gingrich et al., 2000) and other prosocial behaviors (Walsh et al., 2023). Here, RNAseq identified additional genes enriched in the NAc and AI of openers, providing targets for future investigation. Analyzing TFBM in promoters of DEGs revealed that CREB target genes were significantly downregulated in the NAc of the openers group. CREB is upregulated in the NAc following stress and is associated with depressive-like symptoms and dysregulation in motivated behavior (Barrot et al., 2002; Carlezon et al., 2005; Muschamp and Carlezon, 2013; Manning et al., 2017). Thus, CREB downregulation in openers might suggest that helping behavior is protective against a depressive-like phenotype. Additionally, KROX and AP1 activity were elevated in the NAc of openers. As these factors represent immediate-early gene activity, these findings are congruent with the increased levels of c-Fos in openers. Overall, we found that gene expression and transcription factor changes in the NAc, rather than the AI, were related to helping behavior, and OXT signaling in the NAc is a potential target for future manipulations aimed at increasing helping behavior.

Using an in-house open-source freeware, Brainways (Kantor and Bartal, 2023), we identified increased activity in the previously outlined prosocial brain network (Ben-Ami Bartal et al., 2021) for openers compared to non-openers, including in the NAc, AI, OFC, and sensory regions. This finding adds validity to our previous observations of increased NAc activity for trapped ingroup members compared to outgroup members, and further suggests that NAc activity in the HBT reflects affiliation.

The OFC, a region known for its role in goal-directed, value-based and effort-related responding (Wallis, 2007; Münster and Hauber, 2018; Rudebeck and Rich, 2018; Woon et al., 2020), has consistently arisen as a key region active in the HBT. While prior work found the medial OFC to be uniquely active in rats tested with ingroup relative to outgroup members (Ben-Ami Bartal et al., 2021), here, elevated c-Fos was observed across all OFC subregions in openers tested with the same strain, providing support for the involvement of this region in helping behavior, and pointing to valuation processes being involved in helping.

Sensory and insular cortex regions also showed heightened c-Fos activity in openers relative to non-openers. This difference may indicate heightened responsivity to the trapped cagemate, which is associated with an increased likelihood of helping. While past work considered activation of these regions to be common across all conditions of the HBT (regardless of group identity) (Ben-Ami Bartal et al., 2021) here we clearly observe different sensory and insula activity for the same social condition. This suggests that both sensory and insular processing differences are related to the motivational state of the free rat rather than the biological identity of the trapped rat. Lateral and medial ventral thalamus regions were also more active in openers; these regions act as thalamic relay nuclei, conveying sensory information to the insula (Rogers-Carter and Christianson, 2019), providing a connection point for sensory and insular activity in openers. Given the observed differences in social dynamics in openers vs. non-openers, it is possible that neural activity of these regions can be influenced by other parameters, including the relationship between the two animals.

Analysis of functional connectivity in openers identified high connectivity between the ACC, OFC, NAc and PRL, replicating previous reports (Ben-Ami Bartal et al., 2021; Breton et al., 2022). The ACC, a region associated with empathy-like behavior in humans and rodents (Decety et al., 2016), appeared as a major hub within this network, supporting prior work suggesting that connectivity between the ACC and target regions, not just overall ACC activity, is increased in helping conditions.

The current study has several limitations. Experiment 1 was conducted with Wistars, and Experiment 2 with SDs. The observed changes in gene expression should be tested with a SD strain in the future, to ensure that findings translate across multiple rat strains. Additionally, although female rats were included in Experiment 1, Experiment 2 used only males. It remains possible that, though behavioral manifestations of helping were similar across sexes, the biological mechanisms driving this behavior are distinct. Here, gene expression changes were similar across sexes, yet c-Fos data was only collected from males. Future work will need to analyze neural activity in both sexes; this is especially critical as others have reported neural but not behavioral differences in helping behavior in male and female rats (Cox et al., 2024). Another limitation of this study is the impossibility of knowing whether differences in gene expression occurred prior to helping, or due to experience in the HBT. Future work can test for a causal role of *Oxtr* through genetic manipulation or knockdown, and should examine *Oxtr* expression across additional brain regions. Finally, it is impossible to dissociate trait sociability from dyadic interactions, and thus, future studies should aim to control for trait sociability prior to testing to assess the impact of dyadic dynamics on helping behavior.

In sum, these data suggest that social affiliation plays a critical role in variability in prosocial helping behavior. While it is well supported that mammals tend to help those that are familiar/ ingroup members, these findings suggest that relationship strength is predictive of helping even within the ingroup. This work expands and further cements the critical role of the NAc in helping behavior and implicates NAc *Oxtr* levels in contributing towards prosocial helping. Together, this work holds important implications for understanding differences in prosocial motivations and provides targetable mechanisms for future study.

## Conflict of interest statement

The authors declare no competing financial interests

## Acknowledgements

The authors thank Avital Horowitz and Yoni Loterstein for their help with data collection and manual coding for Experiment 1. Parts of some figures were generated with BioRender.com.

## Funding

Israel Science Foundation (IBB), Azrieli Foundation (IBB), Milner Foundation (ET). RH is supported by a President’s Fellowship for Excellent Doctoral Students, Bar-Ilan University.

## Notes

### Competing Interest Statement

The authors have declared no competing interest.

