## Supplemental Material for "Helping behavior is associated with increased affiliative behavior, activation of the prosocial brain network and elevated oxytocin receptor expression in the nucleus accumbens"

***Supplemental Figures***

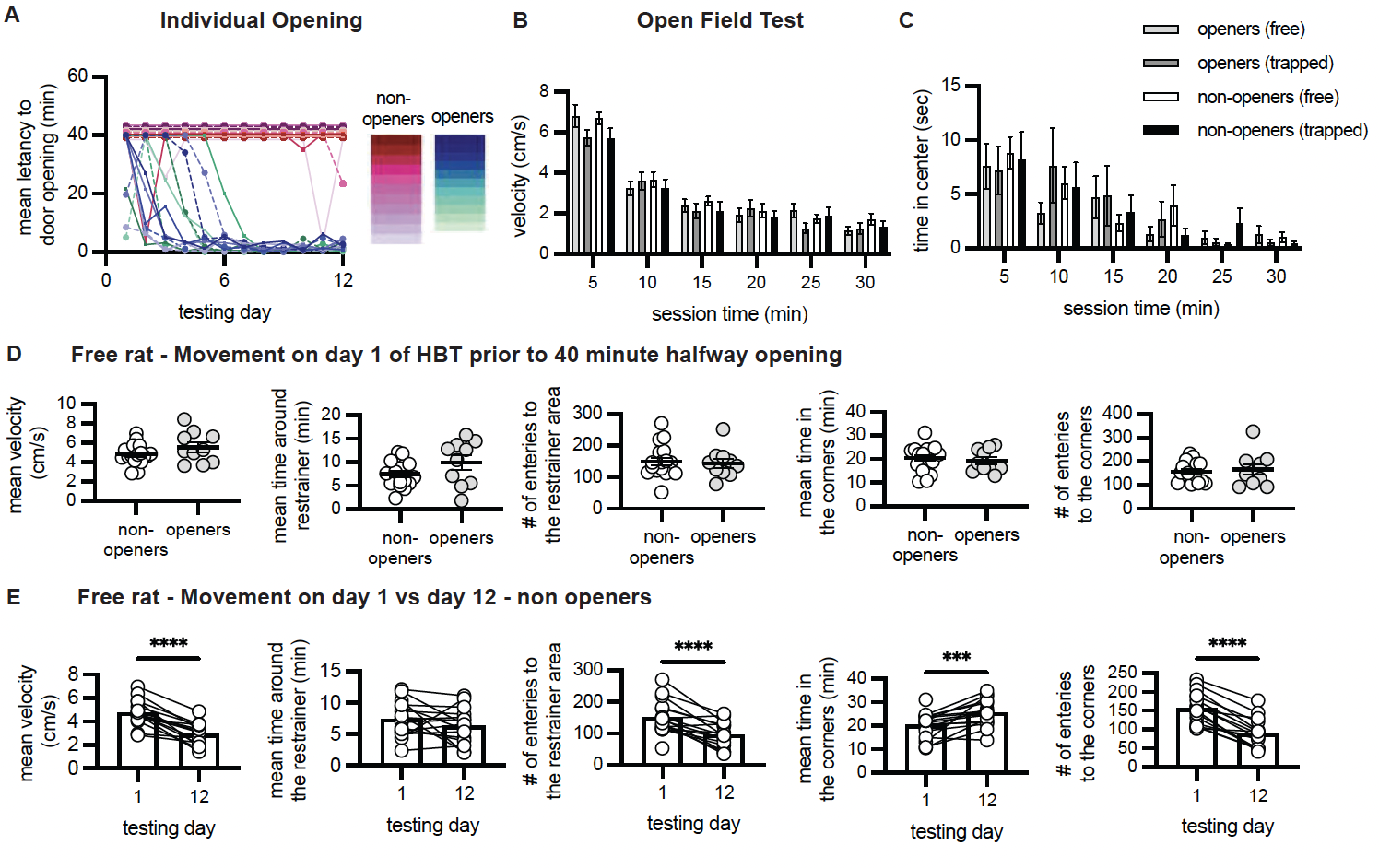

**Figure S1.** A. Mean latency to door-opening across testing sessions for each individual animal. B-C. Velocity (B) and time spent in the center (C) of the 30-minute open field test did not differ across any condition. D. Movement patterns for the free rat on day 1 of helping (including velocity, time and number of entries to the restrainer area and time and number of entries to the corner) did not differ between non-openers and openers. E. For non-openers, movement patterns were altered by day 12, with reduced velocity and entries into the restrainer zone, and more entries and time spent in the corners.

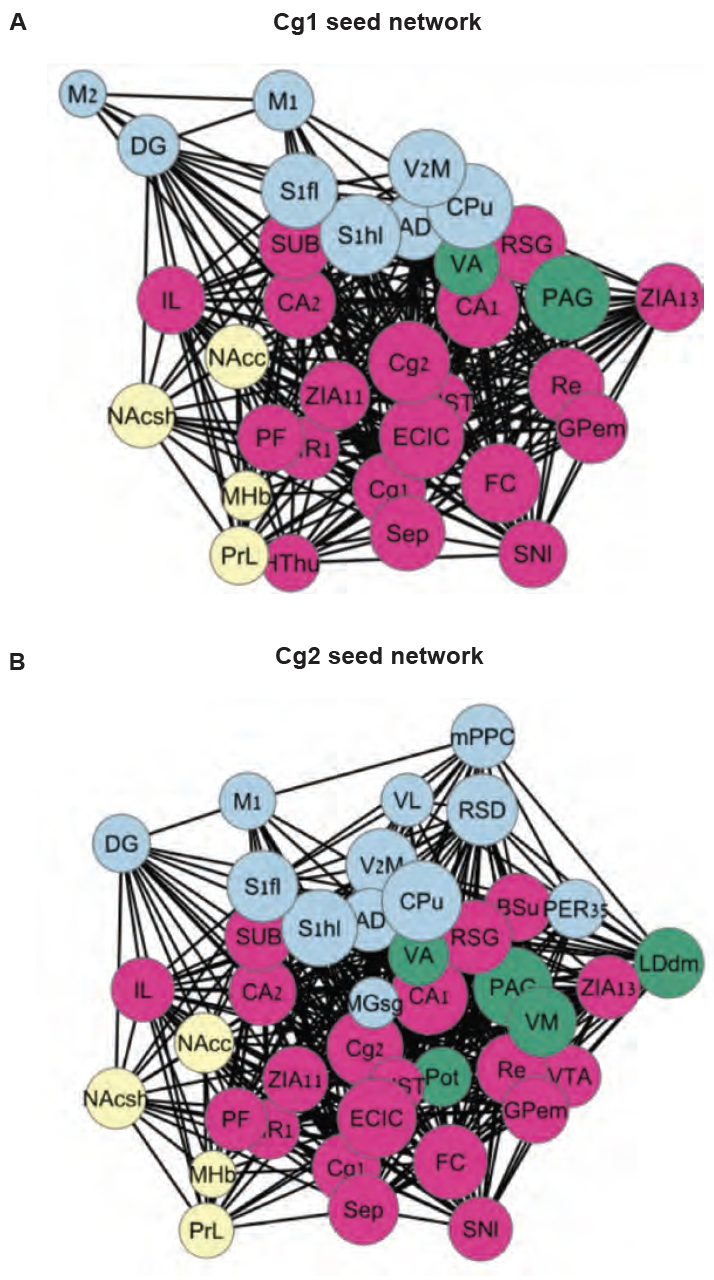

**Figure S2.** Seed network analyses based off of the Cg1 (A) and Cg2 (B) as the primary seed.

***Supplemental Tables***

|  | NaC | | | AI | |
| --- | --- | --- | --- | --- | --- |
| Gene | **Log2 Fold-change** | | **P.val** | **Log2 Fold-change** | **P.val** |
| *Oxtr* | **2.60** | **0.01** | | 0.95 | 0.46 |
| *Drd1* | 0.96 | 0.34 | | 0.93 | 0.73 |
| *Drd2* | 0.96 | 0.38 | | 0.91 | 0.77 |
| *CrhR1* | 1.44 | 0.13 | | 0.92 | 0.80 |
| *Fosl1* | **2.07** | **0.02** | | 0.85 | 0.39 |
| *Nr3c1* | 0.96 | 0.12 | | **1.19** | **0.03** |

**Table S1.** List of *a priori* genes analyzed in Figure 3.

| **NAC** | | | | | |
| --- | --- | --- | --- | --- | --- |
| **TFBM (up)** | **Mean log ratio** | **p value** | **TFBM (down)** | **Mean log ratio** | **p value** |
| V$KROX_Q6 | 1.6558 | 0.0001 | V$ATF_B | -1.047 | 0.0015 |
| V$ZNF219_01 | 1.3623 | 0.0021 | V$ATF_01 | -1.0227 | 0.0001 |
| V$HOXC8_01 | 1.1637 | 0.0071 | V$CREB_02 | -0.9969 | 0.0052 |
| V$POU1F1_Q6 | 1.1528 | 0.0001 | V$ZIC1_01 | -0.9879 | 0.0001 |
| V$TBX5_Q5 | 1.1428 | 0.0064 | V$CREBP1_Q2 | -0.9836 | 0.0051 |
| V$CNOT3_01 | 1.0398 | 0.0068 | V$CREBATF_Q6 | -0.9534 | 0 |
| V$ATATA_B | 1.0239 | 0.0131 | V$E2F1DP1RB_01 | -0.9253 | 0.0151 |
| V$HOXD3_01 | 1.0119 | 0.018 | V$CREB_Q4 | -0.8967 | 0.0064 |
| V$IPF1_03 | 0.9405 | 0.0004 | V$CREB_01 | -0.8531 | 0.0001 |
| V$SP4_Q5 | 0.9071 | 0 | V$GC_01 | -0.8217 | 0.0064 |
| V$SP1_Q2_01 | 0.854 | 0 | V$MYC_Q2 | -0.7874 | 0.0006 |
| V$SP1SP3_Q4 | 0.8533 | 0.0007 | V$USF_01 | -0.7605 | 0.0023 |
| V$SOX9_B1 | 0.7942 | 0.0007 | V$E2F1_Q4 | -0.7597 | 0.0169 |
| V$OCT1_08 | 0.7706 | 0.135 | V$UF1H3BETA_Q6 | -0.7561 | 0.1084 |
| V$NF1_Q6_01 | 0.7665 | 0.0011 | V$HOXA7_01 | -0.7419 | 0.0001 |
| V$AP1_Q4 | 0.7551 | 0 | V$CLOCKBMAL_Q6 | -0.7369 | 0.0009 |
| V$LHX3_01 | 0.7325 | 0.0583 | V$MYCMAX_02 | -0.7239 | 0.0056 |
| V$LEF1TCF1_Q4 | 0.7318 | 0.0007 | V$HNF4_Q6 | -0.7045 | 0.0005 |
| V$PAX7_01 | 0.6917 | 0.0002 | V$FOXM1_01 | -0.7017 | 0.0001 |
| V$TATA_C | 0.649 | 0.0019 | V$MYCMAX_03 | -0.6929 | 0.0031 |
| V$CKROX_Q2 | 0.6362 | 0 | V$SP1_Q6 | -0.687 | 0.0187 |
| V$AP1FJ_Q2 | 0.6354 | 0 | V$CREB_Q4_01 | -0.6867 | 0.001 |
| V$GATA3_02 | 0.5994 | 0.0072 | V$E2F1_Q6 | -0.6842 | 0.0326 |
| V$KAISO_01 | 0.5993 | 0.0538 | V$AP3_Q6 | -0.6782 | 0 |
| **AI** | | | | | |
| **TFBM (up)** | **Mean log ratio** | **p value** | **TFBM (down)** | **Mean log ratio** | **p value** |
| V.HOXA13_02 | 0.87758961 | 0.0007 | V.HDX_01 | -1.33959863 | 0 |
| V.E2F4DP1_01 | 0.77849003 | 0.0109 | V.ICSBP_Q6 | -0.88096488 | 0 |
| V.STAT5A_01 | 0.74757366 | 0 | V.OCT_C | -0.80824998 | 0.0001 |
| V.IPF1_03 | 0.71024876 | 0.0486 | V.GADP_01 | -0.75351258 | 0 |
| V.MYCMAX_02 | 0.6993649 | 0.0004 | V.UF1H3BETA_Q6 | -0.69829853 | 0 |
| V.CHX10_01 | 0.68623627 | 0.0249 | V.TBX5_Q5 | -0.68804172 | 0 |
| V.E2F_02 | 0.58791368 | 0.0303 | V.MEF2_05 | -0.68032805 | 0.0004 |
| V.E2F1DP1RB_01 | 0.58210072 | 0.0218 | V.HMX1_01 | -0.61891849 | 0.0081 |
| V.P53_02 | 0.56891101 | 0 | V.CP2_02 | -0.61599323 | 0.0004 |
| V.STAT5B_01 | 0.53841732 | 0.0063 | V.NKX3A_01 | -0.58992233 | 0.0003 |
| V.ETS1_B | 0.52892685 | 0.0006 | V.RBPJK_01 | -0.57797465 | 0.0008 |
| V.STAT_01 | 0.5174472 | 0.0003 | V.LDSPOLYA_B | -0.57229938 | 0.0003 |
| V.NFY_Q6 | 0.51671654 | 0.0077 | V.HIF1_Q5 | -0.53736018 | 0.0005 |
| V.PROP1_01 | 0.51056809 | 0.0228 | V.PAX_Q6 | -0.53276098 | 0.0068 |
| V.NFY_01 | 0.50975479 | 0.0017 | V.GR_Q6 | -0.51029954 | 0 |
| V.HNF6_Q6 | 0.49147408 | 0.0324 | V.MEF2_Q6_01 | -0.49734942 | 0.0137 |
| V.CAAT_01 | 0.4839906 | 0.0027 | V.BRN2_01 | -0.47988718 | 0 |
| V.TITF1_Q3 | 0.45442331 | 0.0523 | V.T3R_01 | -0.45543744 | 0.083 |
| V.CETS1P54_03 | 0.45141393 | 0.0126 | V.ETS2_B | -0.45183229 | 0.0003 |
| V.AML_Q6 | 0.44855058 | 0.0106 | V.ARNT_02 | -0.44485623 | 0.0291 |
| V.E2F1DP1_01 | 0.43528292 | 0.0473 | V.VMYB_01 | -0.43467575 | 0 |
| V.NRF1_Q6 | 0.42990587 | 0.0469 | V.MYOD_Q6_01 | -0.4124555 | 0.0026 |
| V.GATA3_03 | 0.4276832 | 0.0089 | V.NANOG_01 | -0.41002324 | 0.0155 |
| V.NFY_Q6_01 | 0.37870407 | 0.0088 | V.MYOD_01 | -0.40738891 | 0.0033 |
| V.HEB_Q6 | 0.36202429 | 0 | V.MAF_Q6_01 | -0.3975848 | 0.0413 |

**Table S2.** Top 25 statistically significantly up-regulated and down-regulated transcription factor binding motifs (TFBMs) in promoters of differentially expressed genes within the nucleus accumbens (NAc) and anterior insula (AI). Positive values indicate increased expression in openers relative to non-openers.

| Acronym | Name | Region # | Category |
| --- | --- | --- | --- |
| NLOT | Nucleus of the lateral olfactory tract | 1 | Sensory |
| OBu | Olfactory bulb, unspecified | 2 | Sensory |
| PIR1 | Piriform cortex, layer 1 | 3 | Sensory |
| PIR2 | Piriform cortex, layer 2 | 4 | Sensory |
| PIR3 | Piriform cortex, layer 3 | 5 | Sensory |
| Au1 | Primary auditory area | 6 | Sensory |
| Au2d | Secondary auditory area, dorsal part | 7 | Sensory |
| Au2v | Secondary auditory area, ventral part | 8 | Sensory |
| V1 | Primary visual area | 9 | Sensory |
| V2L | Secondary visual area, lateral part | 10 | Sensory |
| V2M | Secondary visual area, medial part | 11 | Sensory |
| S1bf | Primary somatosensory area, barrel field | 12 | Sensory |
| S1dz | Primary somatosensory area, dysgranular zone | 13 | Sensory |
| S1f | Primary somatosensory area, face representation | 14 | Sensory |
| S1fl | Primary somatosensory area, forelimb representation | 15 | Sensory |
| S1hl | Primary somatosensory area, hindlimb representation | 16 | Sensory |
| S1tr | Primary somatosensory area, trunk representation | 17 | Sensory |
| S2 | Secondary somatosensory area | 18 | Sensory |
| M1 | Primary motor area | 19 | Sensory |
| M2 | Secondary motor area | 20 | Sensory |
| Endo | Endopiriform nucleus | 21 | Association |
| lPPC | Parietal association cortex, lateral area | 22 | Association |
| mPPC | Parietal association cortex, medial area | 23 | Association |
| PtP | Parietal association cortex, posterior area | 24 | Association |
| TeA | Temporal association cortex | 25 | Association |
| Fr3 | Frontal association area 3 | 26 | Association |
| Cg1 | Cingulate area 1 | 27 | Frontal cortex |
| Cg2 | Cingulate area 2 | 28 | Frontal cortex |
| IL | Infralimbic area | 29 | Frontal cortex |
| PrL | Prelimbic area | 30 | Frontal cortex |
| RSD | Retrosplenial dysgranular area | 31 | Frontal cortex |
| RSG | Retrosplenial granular area | 32 | Frontal cortex |
| LO | Lateral orbital area | 33 | Frontal cortex |
| MO | Medial orbital area | 34 | Frontal cortex |
| VO | Ventral orbital area | 35 | Frontal cortex |
| VLO | Ventrolateral orbital area | 36 | Frontal cortex |
| DLO | Dorsolateral orbital area | 37 | Frontal cortex |
| AId | Agranular insular cortex dorsal area | 38 | Insula |
| AIp | Agranular insular cortex, posterior area | 39 | Insula |
| AIv | Agranular insular cortex, ventral area | 40 | Insula |
| CLA | Claustrum | 41 | Insula |
| DI | Dysgranular insular cortex | 42 | Insula |
| GI | Granular insular cortex | 43 | Insula |
| Amu | Amygdaloid area, unspecified | 44 | Amygdala |
| BNST | Bed nucleus of the stria terminalis | 45 | Amygdala |
| CA1 | Cornu ammonis 1 | 46 | Hippocampal formation |
| CA2 | Cornu ammonis 2 | 47 | Hippocampal formation |
| CA3 | Cornu ammonis 3 | 48 | Hippocampal formation |
| DG | Dentate gyrus | 49 | Hippocampal formation |
| FC | Fasciola cinereum | 50 | Hippocampal formation |
| LEC | Lateral entorhinal cortex | 51 | Hippocampal formation |
| PaS | Parasubiculum | 52 | Hippocampal formation |
| PER35 | Perirhinal area 35 | 53 | Hippocampal formation |
| PER36 | Perirhinal area 36 | 54 | Hippocampal formation |
| PrS | Presubiculum | 55 | Hippocampal formation |
| SUB | Subiculum | 56 | Hippocampal formation |
| CPu | Caudate putamen | 57 | Striatum |
| NAcc | Nucleus accumbens, core | 58 | Striatum |
| NAcsh | Nucleus accumbens, shell | 59 | Striatum |
| Sep | Septal region | 60 | Striatum |
| VP | Ventral pallidum | 61 | Striatum |
| VSRu | Ventral striatal region, unspecified | 62 | Striatum |
| BFRu | Basal forebrain region, unspecified | 63 | Basal Ganglia |
| EP | Entopeduncular nucleus | 64 | Basal Ganglia |
| GPel | Globus pallidus external, lateral part | 65 | Basal Ganglia |
| GPem | Globus pallidus external, medial part | 66 | Basal Ganglia |
| RTa | Reticular (pre)thalamic nucleus, auditory segment | 67 | Prethalamus |
| RTu | Reticular (pre)thalamic nucleus, unspecified | 68 | Prethalamus |
| LHb | Lateral habenular nucleus | 69 | Epithalamus |
| MHb | Medial habenular nucleus | 70 | Epithalamus |
| AD | Anterodorsal thalamic nucleus | 71 | Thalamus |
| AM | Anteromedial thalamic nucleus | 72 | Thalamus |
| AVdm | Anteroventral thalamic nucleus, dorsomedial part | 73 | Thalamus |
| AVvl | Anteroventral thalamic nucleus, ventrolateral part | 74 | Thalamus |
| IAM | Interanteromedial thalamic nucleus | 75 | Thalamus |
| IMD | Intermediodorsal thalamic nucleus | 76 | Thalamus |
| PIL | Posterior intralaminar nucleus | 77 | Thalamus |
| PT | Parataenial thalamic nucleus | 78 | Thalamus |
| PV | Paraventricular thalamic nuclei (anterior and posterior) | 79 | Thalamus |
| SPF | Subparafascicular nucleus | 80 | Thalamus |
| CL | Central lateral thalamic nucleus | 81 | Thalamus |
| CM | Central medial thalamic nucleus | 82 | Thalamus |
| Eth | Ethmoid-Limitans nucleus | 83 | Thalamus |
| PCN | Paracentral thalamic nucleus | 84 | Thalamus |
| PF | Parafascicular thalamic nucleus | 85 | Thalamus |
| DLG | Dorsal lateral geniculate nucleus | 86 | Thalamus |
| LDdm | Laterodorsal thalamic nucleus, dorsomedial part | 87 | Thalamus |
| LDvl | Laterodorsal thalamic nucleus, ventrolateral part | 88 | Thalamus |
| LPl | Lateral posterior thalamic nucleus, lateral part | 89 | Thalamus |
| LPmc | Lateral posterior thalamic nucleus, mediocaudal part | 90 | Thalamus |
| LPmr | Lateral posterior thalamic nucleus, mediorostral part | 91 | Thalamus |
| MDc | Mediodorsal thalamic nucleus, central part | 92 | Thalamus |
| MDl | Mediodorsal thalamic nucleus, lateral part | 93 | Thalamus |
| MDm | Mediodorsal thalamic nucleus, medial part | 94 | Thalamus |
| MGd | Medial geniculate body, dorsal division | 95 | Thalamus |
| MGm | Medial geniculate body, medial division | 96 | Thalamus |
| MGmz | Medial geniculate body, marginal zone | 97 | Thalamus |
| MGsg | Medial geniculate body, suprageniculate nucleus | 98 | Thalamus |
| MGv | Medial geniculate body, ventral division | 99 | Thalamus |
| SubG | Subgeniculate nucleus | 100 | Thalamus |
| IGL | Intergeniculate leaflet | 101 | Thalamus |
| PrG | Pregeniculate nucleus | 102 | Thalamus |
| Po | Posterior thalamic nucleus | 103 | Thalamus |
| Pot | Posterior thalamic nuclear group, triangular part | 104 | Thalamus |
| Re | Reuniens thalamic nucleus | 105 | Thalamus |
| Rh | Rhomboid thalamic nucleus | 106 | Thalamus |
| Xi | Xiphoid thalamic nucleus | 107 | Thalamus |
| Ang | Angular thalamic nucleus | 108 | Thalamus |
| SMT | Submedius thalamic nucleus | 109 | Thalamus |
| VA | Ventral anterior thalamic nucleus | 110 | Thalamus |
| VL | Ventrolateral thalamic nucleus | 111 | Thalamus |
| VM | Ventromedial thalamic nucleus | 112 | Thalamus |
| VPL | Ventral posterolateral thalamic nucleus | 113 | Thalamus |
| VPM | Ventral posteromedial thalamic nucleus | 114 | Thalamus |
| VPpc | Ventral posterior nucleus of the thalamus, parvicellular part | 115 | Thalamus |
| STh | Subthalamic nucleus | 116 | Thalamus |
| ZIA11 | Zona incerta, A11 dopamine cells | 117 | Subthalamus |
| ZIA13 | Zona incerta, A13 dopamine cells | 118 | Subthalamus |
| ZIc | Zona incerta, caudal part | 119 | Subthalamus |
| ZId | Zona incerta, dorsal part | 120 | Subthalamus |
| ZIr | Zona incerta, rostral part | 121 | Subthalamus |
| ZIv | Zona incerta, ventral part | 122 | Subthalamus |
| FoF | Fields of Forel | 123 | Subthalamus |
| HThu | Hypothalamic region, unspecified | 124 | Hypothalamus |
| ECIC | Inferior colliculus, external cortex | 125 | Midbrain |
| SuD | Deeper layers of the superior colliculus | 126 | Midbrain |
| SuG | Superficial gray layer of the superior colliculus | 127 | Midbrain |
| IP | Interpeduncular nucleus | 128 | Midbrain |
| PAG | Periaqueductal gray | 129 | Midbrain |
| PP | Peripeduncular nucleus | 130 | Midbrain |
| SNc | Substantia nigra, compact part | 131 | Midbrain |
| SNl | Substantia nigra, lateral part | 132 | Midbrain |
| SNr | Substantia nigra, reticular part | 133 | Midbrain |
| VTA | Ventral tegmental area | 134 | Midbrain |
| Pn | Pontine nuclei | 135 | Midbrain |
| PRT | Pretectal region | 136 | Midbrain |
| BSu | Brainstem, unspecified | 137 | Brainstem |

**Table S3.** Detailed list of 137 brain regions analyzed and used in Figures 4-5. Brain region abbreviations and full names, organized by presentation and brain region category.
